# NuclampFISH enables cell sorting based on nuclear RNA expression for chromatin analysis

**DOI:** 10.1101/2024.10.30.619948

**Authors:** Yifang Liu, Yuchen Qiu, Keqing Nian, Sara H. Rouhanifard

## Abstract

Transcriptional bursts refer to periods when RNA polymerase interacts with a DNA locus, leading to active gene transcription. This bursting activity can vary across individual cells, and analyzing the differences in transcription sites can help identify key drivers of gene expression for a specific target. Scaffolding methods based on fluorescence in situ hybridization (FISH) have been widely used to amplify the fluorescent signal of mRNAs and sort cells based on mRNA expression levels. However, these methods are inefficient at targeting nuclear RNA, including transcription sites, due to the probes’ limited accessibility through cellular compartment membranes and crosslinked proteins. Additionally, the required formaldehyde fixation interferes with downstream analysis of chromatin and protein-binding interactions. To address these challenges, a platform that integrates amplified FISH with reversible crosslinkers and allows access to the nucleus is needed. In response, we developed nuclear clampFISH (nuclampFISH). This method amplifies the fluorescent signal of mRNAs using a reversible crosslinker, enabling the sorting of cells based on nuclear RNA expression and compatible with downstream biochemical analysis. This assay demonstrates that transcriptionally active cells have more accessible chromatin for a respective gene. Notably, the tools developed are highly accessible and do not require specialized computation or equipment.

## Introduction

Transcription sites are specific loci where the RNA polymerase binds to the DNA. The presence of a transcription site by methods such as RNA FISH indicates that these genes are transcriptionally active or “bursting.” It is well established in the field that single cells have high variability in the expression level of transcription sites, meaning they are in distinct cellular states^1–4^. However, the specific mechanisms and regulatory machinery behind such variation are less understood. Transcription regulation, specifically bursting, involves the recruitment and release of RNA Pol II at the promoter. When this activity is very high, the expression of bursts can appear homogeneous, but when they are less frequent, it leads to cell-to-cell variability^5^. One of the major challenges is to understand the regulators of bursting.

Current methods for studying transcription sites primarily use RNA PolII chromatin immunoprecipitation coupled with sequencing (ChIP-seq)^6,7^. This method informs on DNA and protein interactions, thus identifying actively transcribed regions. Other methods are based on nascent RNA enrichment and sequencing, which can track newly transcribed RNAs^8^. However, while these methods are genome-wide and provide details about DNA-protein interactions, these are bulk methods that lack single-cell resolution. Single-molecule fluorescence *in situ* hybridization (smFISH) is an imaging-based method for measuring the frequency and duration of transcription sites in single cells by targeting the introns of transcripts of interest^9,10^. It can provide information on spatial localization and absolute quantification of transcripts by detecting single molecules of RNA inside fixed cells. However, this method does not inform RNA-protein interactions, and the smFISH signal is insufficiently bright enough to separate cells based on transcriptional activity using flow cytometry, which would enable biochemical assays. An ideal method for studying transcriptional bursts would be gene-specific, sensitive, and specific enough for flow cytometry, inform on single-cell activity, and have compatibility with biochemical assays for downstream analysis.

Various FISH-based scaffolding methods have been developed to enhance the fluorescent signals produced from RNA-binding oligonucleotide probes. These methods can all achieve high amplification^11–13^ but lose signal integrity in nuclear RNA. One of these methods we developed is click-amplified FISH (clampFISH), which uses a “C” shaped probe to hybridize and form a double helix with the target RNA, followed by ligation using bioorthogonal click chemistry^14,15^. The benefit of this method is that the ligated probes can survive stringent wash steps and achieve exponential amplification, and the backbone sequence can be modified for facile multiplexing. This method has also been recently expanded for faster analysis and higher throughput (clampFISH 2.0)^15^.

We establish nuclampFISH to target nuclear RNA and transcription sites with a reversible crosslinker. We use this amplification strategy to sort cells for the first time based on transcriptional activity. We perform a chromatin accessibility assay to understand the differences between transcriptionally active and inactive cells.

## Results

### ClampFISH and HCR probes are inaccessible to transcription sites

Previous reports have suggested that nuclear accessibility does not influence smFISH probes permeabilization (**Figure. 1a**), but it presents a challenge for FISH-based amplification, such as clampFISH and Hybridization chain reaction (HCR) (**Figure. 1b,c**)^14^. Transcription sites reside in the nucleus and may be detected by the colocalization of exon and intron FISH probes for the same transcript (**Figure. 1d**). To assess the labeling efficiency of molecular scaffold-based amplification strategies for detecting transcription sites, we labeled *EEF2* mRNA with smFISH, clampFISH, and HCR (**FIgure. 1e**). This target has an average of 1.5 transcription sites in most HeLa cells, as detected by the colocalization of smFISH intron and exon probes in the nucleus (**Figure. 1e**). ClampFISH and HCR probes were designed to target the *EEF2* exon sequence. They were counterstained by the smFISH probes that targeted the *EEF2* intron. We found that clampFISH and HCR can reliably detect exon mRNA in cytoplasm, but the signal from the transcription site was lost or not specific. The nuclear exon spots overlaid well with nuclear intron spots (arrow), indicating that smFISH can detect transcription sites specifically (**Figure. 1e**). The distribution of exon FISH spots in clampFISH, HCR, and smFISH overlaps and the median number of the RNA spots counts in these three conditions are consistent: HCR: 250±4.2, clampFISH: 236±8.5, smFISH: 235±7.4 (**Figure. 1f**).

**Figure 1.**
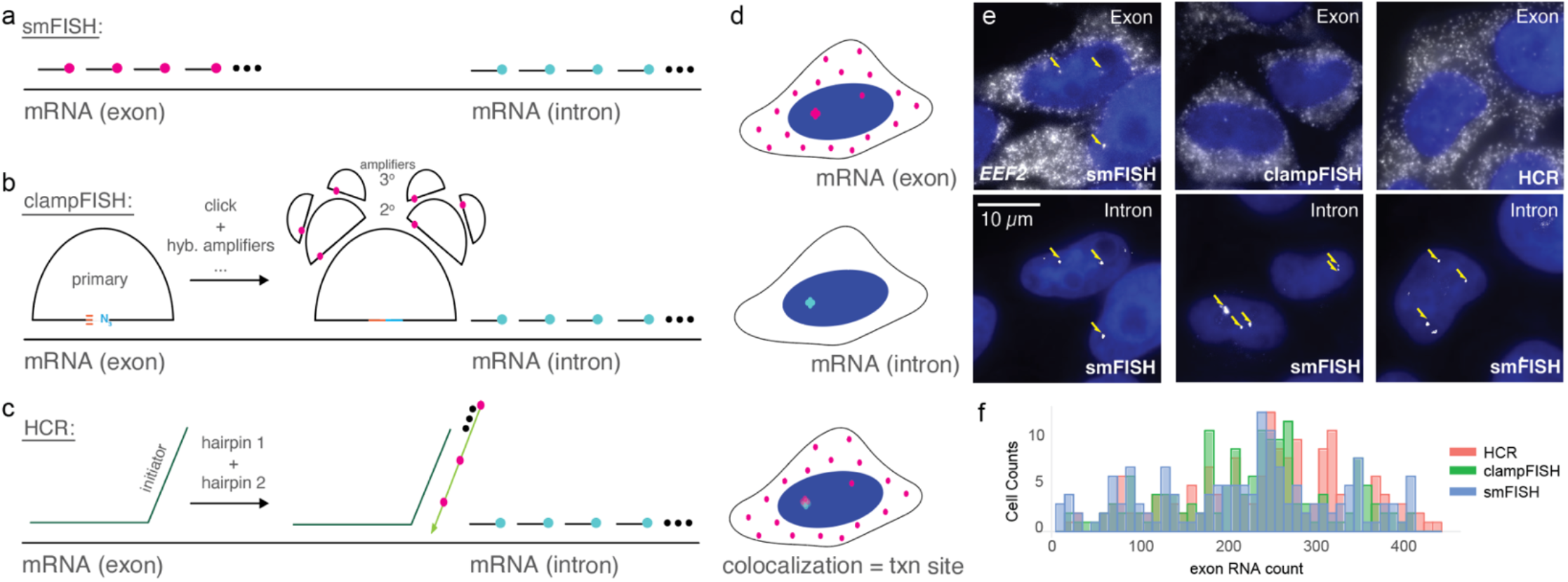
FISH-based methods to detect transcription sites include a. smFISH, b. clampFISH, and c. HCR FISH. d. Overlaid FISH spots in mRNA exon and intron indicate transcription (txn) sites. e. Images of EEF2 exons and smFISH images of EEF2 introns were used to show the colocalization and identify transcription sites. f. Overlaid histogram plot of exon RNA count distribution for clampFISH, HCR FISH, and smFISH for two biological replicates.

### Nuclear isolation and method optimization enable exponential amplification of nuclear RNA

One hypothesis is that FISH amplification strategies are not capable of probing transcription sites because the probe binding is inefficient, and a larger transcriptional burst would not have this problem. To test whether increasing the number of nascent transcripts for our target transcript would enhance the signals to a detectable level, we treated cells with Pladienolide B (Pla B) ^16^. This splicing inhibitor induces the transcription of *EEF2* by blocking the assembling process of spliceosome ^17^. We applied clampFISH probes targeting *EEF2* exon using the original clampFISH protocol^14^ with and without the presence of Pla B. We found that even after Pla B treatment, clampFISH still exhibited a dim signal even though the smFISH signal had a 1.73-fold increase as without Pla B treatment (**Figure. 2a,b, Supplementary Fig. 1**) and minimal 0.78%±0.53% colocalization ratio with the intron smFISH probes ( **Supplementary Fig. 2**).

**Figure 2.**
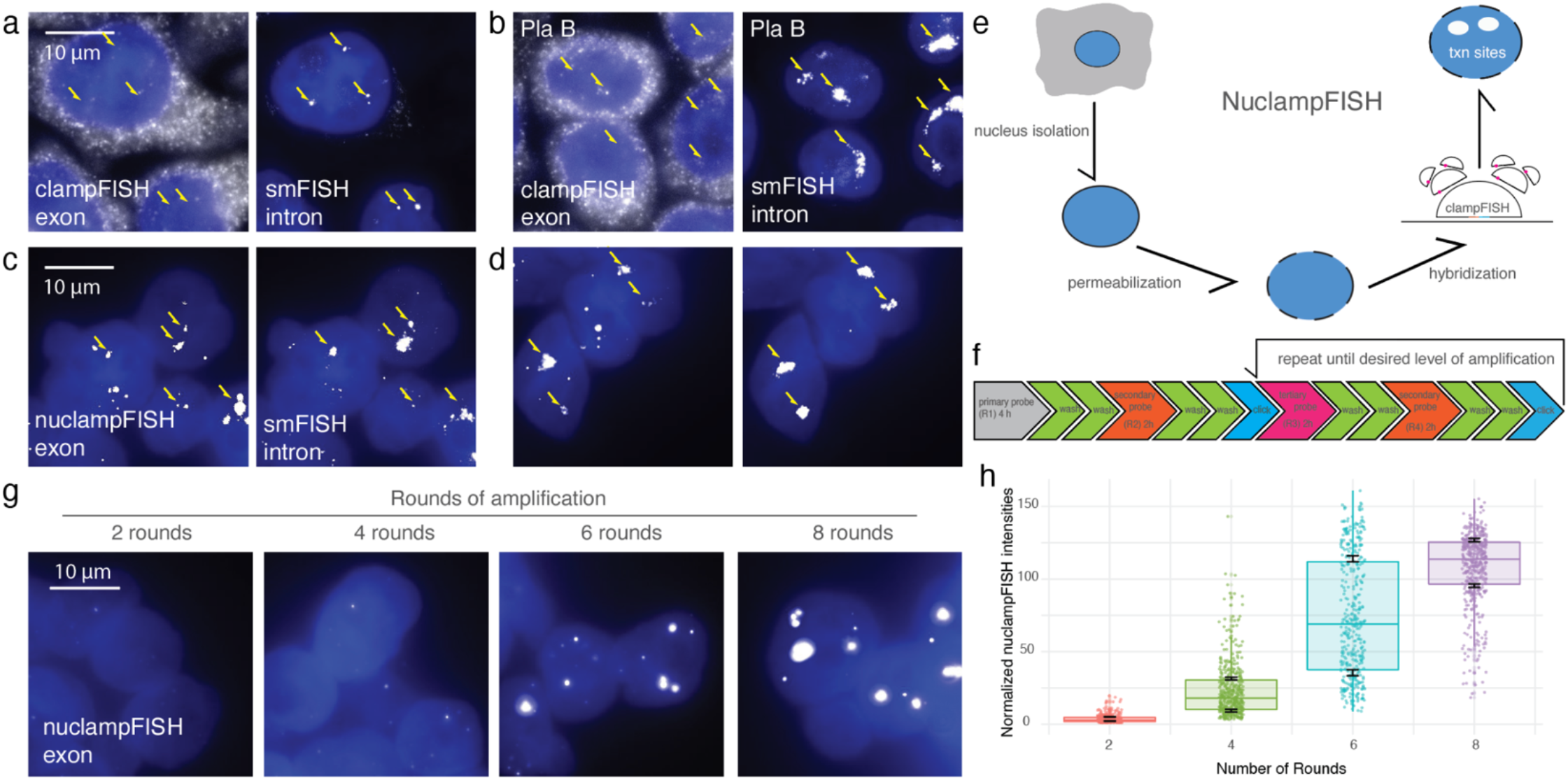
Method development and workflow of nuclampFISH. a. clampFISH images of EEF2 exon and smFISH images of EEF2 intron without Pla B treatment. The arrow indicates the transcription sites. b. clampFISH images of EEF2 exon and smFISH images of EEF2 intron after Pla B treatment. c. NuclampFISH images of EEF2 exon after nucleus isolation and smFISH images of EEF2 intron. d. NuclampFISH images of EEF2 exon after nucleus isolation and modified hybridization parameters, and smFISH images of EEF2 intron. e. Workflow of nuclampFISH methods. f. Timeline of nuclampFISH process. g. Amplification efficiency of nuclampFISH, two rounds, four rounds, six rounds, eight rounds of nuclampFISH images were shown. h. Box and whisker plot of nuclampFISH spots intensities in each round of amplification. n=3 biological replicates.

We next hypothesized that the molecular scaffold size of the clampFISH probe prevents itself from diffusing through the cytoplasm to access the target in the nucleus. We removed the cell membrane and cytoplasm, then applied the clampFISH probes targeting *EEF2* exons with two rounds of amplification (i.e., primary probe and secondary probe). After nuclear isolation, the signal intensities of transcription sites inside the cell nucleus and colocalization ratio with smFISH intron probes were improved significantly (**Figure. 2c, Supplementary Fig. 2**) to 15.7%±4.1%. Next, we optimized the inclusion of surfactant and salt to improve probe binding. Triton X-100 enhances cell permeability, and salt drives probe hybridization with the target mRNA. After adding Triton X-100, we found that the nucleus clampFISH signal intensities and colocalization of clampFISH probes with smFISH intron probes increased to 24.4%±2.1% (**Figure. 2d, Supplementary Fig. 2**). These values further increased by increasing the salt concentration from 2X SSC to 5X SSC (**Figure. 2d, Supplementary Fig. 2**). The same result was observed for another nuclear RNA target, *NEAT1*, a long non-coding RNA (**Supplementary Fig. 3)** and *TMSF1* exons **(Supplementary Fig. 4)**.

Next, we tested the new method of probe delivery that would specifically and exponentially amplify the nuclear signal. We used the same primary probes used for the *EEF2* exon and amplified the signal using fluorescent secondary and tertiary probes with a click reaction performed (**Figure. 2f)**. We performed the nuclampFISH protocol with varying stopping points: two rounds (primary and secondary), four rounds (primary, secondary, tertiary, and secondary again), six rounds, and eight rounds. We observed bright, amplified spots with increasing fluorescent signal as amplification rounds increased (**Figure. 2g, h**). Furthermore, we observed an exponential rate of amplification,1.74-fold per round, indicating that the amplifiers’ binding efficiency is 87.1 % of the theoretical doubling intensity per round. This rate did not slow down even when we reached eight rounds of amplification, indicating that an even brighter signal could be achievable with additional amplification.

### NuclampFISH is compatible with intron targeting and reversible crosslinking

FISH-based methods require fixation and permeabilization before probe hybridization, including those with molecular scaffolding for fluorescence amplification. This fixation is almost universally using formaldehyde. Formaldehyde crosslinking is conventionally used for FISH-based assays^18,19^. However, it can interfere with downstream analysis, such as mass spectrometry, biochemical assays, and chromatin profiling^20,21^. To overcome the challenge of formaldehyde, we tested the compatibility of several alternative crosslinkers with smFISH and clampFISH, including glutaraldehyde, methanol, disuccinimidyl sulfoxide (DSSO), and dithiobis (succinimidyl propionate) (DSP). We found that DSSO and DSP have comparable performance with formaldehyde crosslinking **(Supplementary Fig. 5)**. We chose DSP crosslinking because this is chemically reversible, and DSSO is reversible by mass spectrometry. DSP could be applied to a broader variety of downstream applications. DSP is a crosslinker that can be easily reversed by 10-50 mM dithiothreitol (DTT ) or tris(2-carboxyethyl)phosphine(TCEP) at pH 8.5^22^. We compared the performance of formaldehyde and DSP fixation by targeting the *HIST1H4E* mRNAs using clampFISH exon probes (**Figure. 3a**). We found that the RNA count distribution of DSP crosslinking can be overlaid well with formaldehyde crosslinking **(Figure. 3b)**. We applied the nuclampFISH method to target transcription sites using DSP crosslinking with probes complementary to the *EEF2* intron mRNA sequence. We observed that the nuclampFISH spots colocalized with smFISH spots at transcription sites, indicating that the protocol is compatible with DSP for targeting nuclear RNA while maintaining specificity (**Figure. 3c**).

**Figure 3.**
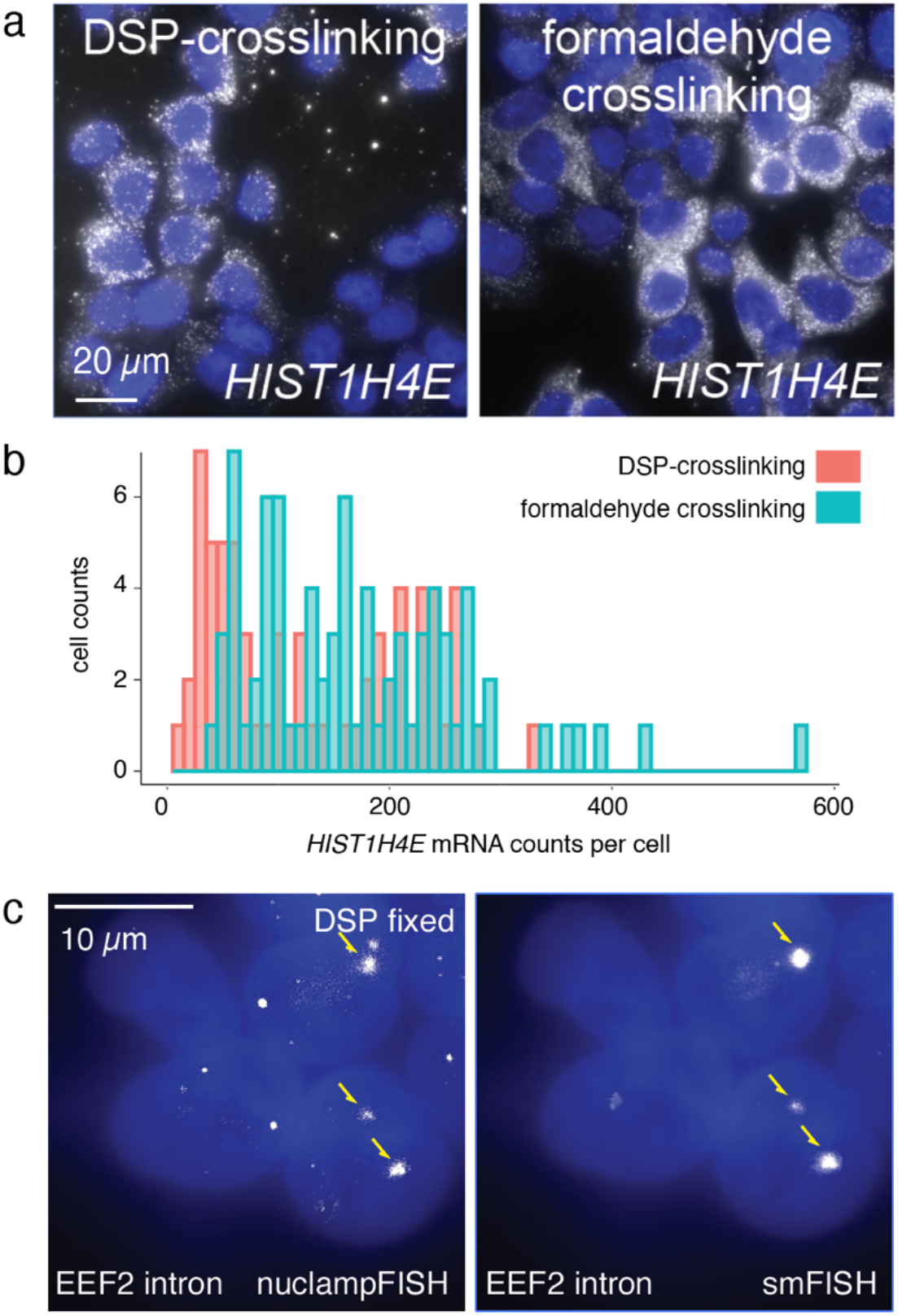
Application of reversible crosslinker to nuclampFISH. a. Photomicrograph of HeLa cells stained for HIST1H4E mRNA using clampFISH after DSP (left) and formaldehyde crosslinking (right). b. Histogram of HIST1H4E mRNA counts per cell in DSP crosslinked HeLa cells versus formaldehyde crosslinked HeLa cells. c. NuclampFISH images of EEF2 intron after nucleus isolation and smFISH images of EEF2 intron.

### NuclampFISH enables cell sorting based on nuclear RNA expression

One primary application is that nuclampFISH enables sorting cells based on nuclear RNA expression by detecting the active transcription sites of a single mRNA. We used nuclampFISH to sort cells based on the expression of actively transcribing *EEF2* (i.e., clampFISH primary probes targeting the *EEF2* intron). Compared to the negative control, which included each round of amplifier probe, the positive group, which was treated with four rounds of amplification, showed almost a complete decade shift (**Figure. 4b**). We set three gates (G1, G2, G3 with increased nuclampFISH signal) to sort cells based on the expression level of transcription sites (**Figure. 4b**). After collecting the cells from each gate, we imaged them using fluorescence microscopy (**Figure. 4c**). We found that the nuclampFISH signal on transcription sites, as shown by microscopy, corresponded to the increasing fluorescent intensity observed by flow cytometry (**Figure. 4c,d**). We extracted the RNA from the sorted cells and conducted RT-qPCR to quantify the levels of *EEF2* intron from each gated population. The RNA counts correlated significantly with nuclampFISH pixel intensities acquired by microscopy after sorting (r=0.93; **Figure. 4d**) and nuclampFISH spots quantity (**Supplementary Fig. 6)**.

**Figure 4.**
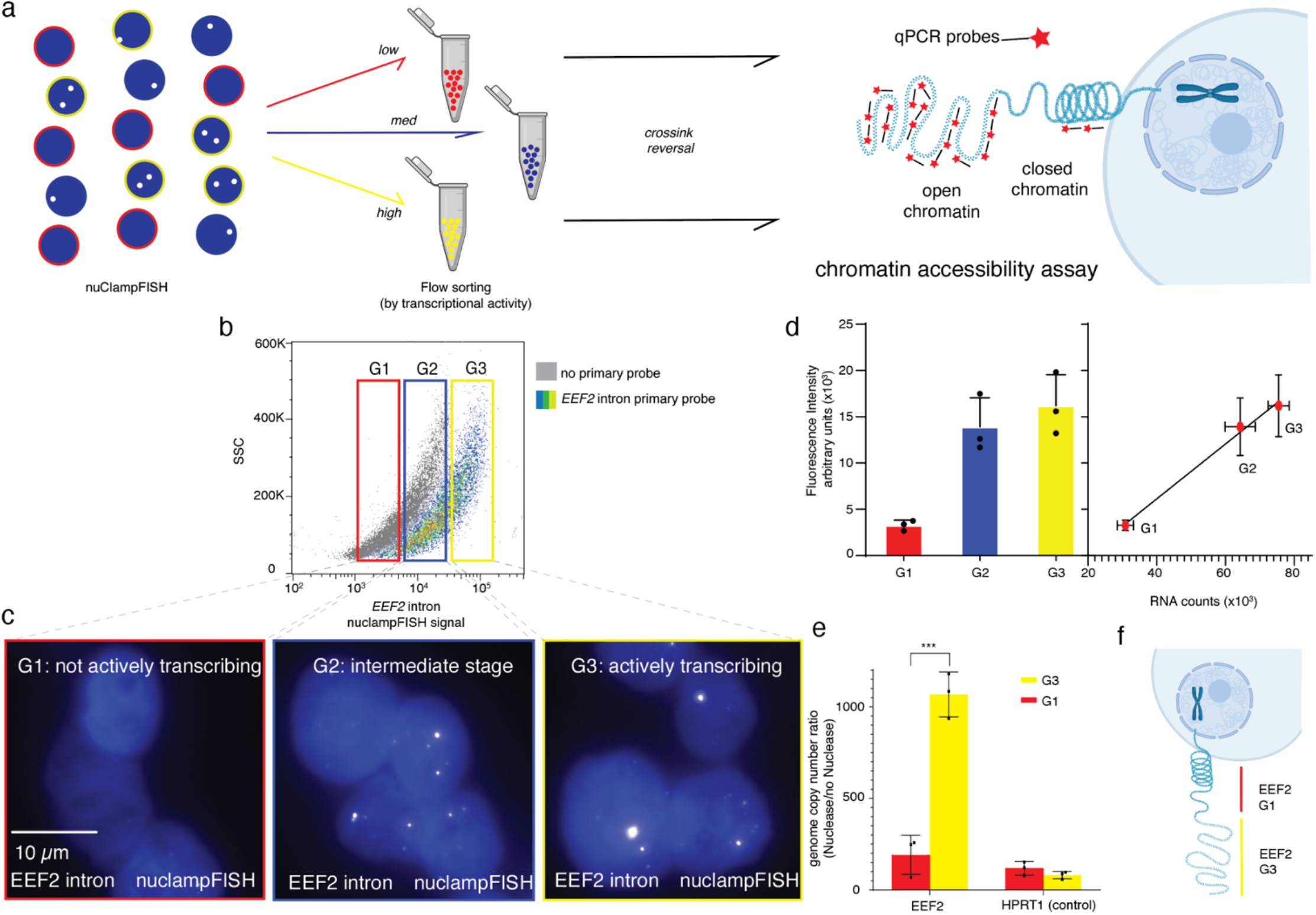
a. Workflow of nuclampFISH, flow sorting, and chromatin accessibility assay. b. A flow cytometry plot (pseudocolored) shows a gating strategy for EEF2 intron mRNA in HeLa cells. Low, medium, and high expressers were gated as G1, G2, and G3, respectively. c. fluorescent micrographs of G1, G2, and G3 cell populations after sorting. d. (left) Bar graph of quantified nuclampFISH signal in each cell population after flow sorting (right) Correlation of RT-pPCR counts with nuclampFISH signal intensities after flow sorting. e. To quantify chromatin accessibility for the G1 and G3 groups, the ratio of qPCR counts of nuclease treatment to no nuclease treatment was measured to determine the quantity of chromatin open level. f. Cartoon modeling chromatin openness for G1 and G3 groups. n=3 biological replicates (mean ± SD).

### NuclampFISH enables cell sorting and crosslink reversal for chromatin accessibility analysis

Chromatin accessibility assays such as DNA sequencing-based methods facilitate characterizing active regulatory elements during active transcription^23,24^. However, these are typically bulk assays that combine heterogeneous cell populations for analysis rather than single-cell assays. Single-cell assays such as smFISH can analyze gene-specific, nascent RNA; however, these assays do not inform DNA accessibility. We sought to combine the single-cell and gene-specific benefits of FISH-based assays with the sensitivity of bulk assays to detect chromatin structure and explore chromatin conformation during active transcription of a given gene.

As a critical first step, we hypothesized that the chromatin accessibility assay would be compatible with DSP-crosslinked cells for which the crosslinking was reversed. Chromatin accessibility assays report on the “openness” of chromatin based on the effectiveness of qPCR in a region of interest. Crosslinking the nucleic acids and proteins surrounding the region of interest can lead to misinterpretation of the results as closed when it could be a crosslinking artifact. As a proof-of-concept, we crosslinked the cells with DSP and compared the *EEF2* chromatin accessibility of the DSP reversal group with the no DSP reversal group; the reversal group exhibited a 3.16-fold increase in signal **(Supplementary Fig. 7**), indicating that crosslink reversal is compatible with the chromatin accessibility assay.

Next, we DSP crosslinked HeLa cells, labeled *EEF2* introns using nuclampFISH with four rounds of amplification, and collected the sorted nucleus from the G1 and G3 groups. Crosslinking was reversed from the G1 and G3 groups, followed by chromatin accessibility analysis for the genomic region harboring the *EEF2* intron. We observed that the G3 group (i.e., the group with a higher level of “burstiness” for *EEF2* mRNA) had a 5.59-fold increase in accessibility compared to the G1 group (i.e., the group with limited “burstiness” for *EEF2* mRNA; *p*<0.0001; **Figure. 4e**). As a critical control, we selected a housekeeping gene, *HPRT1*, and performed the chromatin accessibility assay on the G1 and G3 groups selected for *EEF2. HPRT1* did not show a significant difference in chromatin accessibility (**Figure. 4e**). These results demonstrate that “bursty” transcription corresponds with more accessible chromatin at the gene of interest **(Figure. 4f)**.

## Discussion

Here, we established nuclampFISH, an amplified FISH method for targeting nuclear RNA and transcription sites. This method achieves the goal of specific detection of nuclear RNA (including transcription sites) and amplifying the FISH signal of nuclear RNA (including transcription sites). Based on the specific and amplified nuclampFISH signal, we offer a platform to separate cells according to the expression level of transcription sites for downstream analysis to understand transcription better. We combine the FISH detection and chromatin analysis with a chemical reversible crosslinker DSP.

We demonstrated that, when compared to clampFISH and HCR FISH for transcription sites, nuclampFISH can significantly increase the signal and achieve specific detection. Moreover, nuclampFISH can maintain the exponential amplification capacity of clampFISH. These methods can detect multiple nuclear RNAs, such as the lncRNA *NEAT1*. However, the total fluorescence produced for amplification strategies targeting *NEAT1* is lower than expected based on the amplification increase for cytoplasmic RNAs, suggesting that the labeling efficiency is also lower for nuclear RNAs.

Next, we sorted the cells based on the expression level of transcriptional bursts. The accuracy of the separation was demonstrated by both RT-qPCR and imaging. After sorting and collecting nuclei based on expression levels, we reversed the crosslinking and performed a chromatin accessibility assay for the gene of interest. We observed that transcriptionally active cells had more open levels of chromatin than the transcription inactive cells. This provides direct evidence that open chromatin regions enable the binding of transcription factors and other regulatory elements to drive the transcription of a given gene. In the stochastic bursting model, a simplistic model fits kinetic information in a two-state system where the promoter alternates between an ON and OFF position^25^. This model has been expanded to include a third, ‘refractory’ promoter state where the promoter is bound but resistant to activation (i.e., no burst). Our data shows that the cells exhibiting no bursting have significantly less “open chromatin” than the bursting cells, with levels comparable to that of negative control, *HPRT1*. This supports the two-state model.

Importantly, our established platform combines single-cell analysis with bulk-cell sensitivity to move beyond an RNA-centric view of transcription and include the interacting factors of chromatin and protein in the analysis. The sorting and crosslinking reversal steps will enable future downstream analysis such as chromatin capture, mass spectrometry (MS), and biochemical assays such as western blot and immunoprecipitation.

In summary, we report developing a method for the specific detection of nuclear RNA and transcription sites. This assay broadens the application of clampFISH for transcription sites by amplifying specific FISH signals, thus enabling the separation of the cells for downstream analysis, including chromatin analysis, proteomics, and transcriptional profiling.

## Methods

### Tissue culture conditions

HeLa cells were handled according to the manufacturer’s protocol and maintained in high-glucose DMEM with GlutaMax (FisherSci cat# 10566016), 10% FBS, and 1% Pen-Strep (FisherSci cat# BW17-602E). Cell lines tested PCR-negative for mycoplasma contamination (ATCC cat# 30-1012K).

### Drug treatment for splicing inhibition

Cells were treated with 1 µM Pladienolide B (Tocris Biosciences #6070500U) for 4 h to achieve splicing inhibition as described by previous work ^26^.

### smFISH

The smFISH detection was performed as previously described^19^. Sequences for smFISH probes can be found in **Supplementary Table 1**

### HCR FISH

HCR FISH was conducted using the HCR™ RNA-FISH Bundle from Molecular Instruments. We followed the manufacturer’s protocol ^11,27^.

### NuclampFISH protocol

#### Nucleus extraction

Minute^™^ Single Nucleus Isolation Kit for Tissues/Cells (Invent Biotechnologies cat# SN-047) was used to extract nuclei from whole cells. 5 × 10 ^6^-10 ^7^ cells were collected and washed by pre-cold 1X PBS once. 200 µL lysis buffer was added to the tube, and the cells were ground with the pestle provided 20-30 times. Another 400 µL lysis buffer was added to the tube and incubated at 4°C for 10 min. After incubation, all cell lysate was transferred into a filter in a collection tube. The nuclei pellet was obtained by centrifuge at 600 X g for 5 min. The supernatant was removed carefully and discarded. The pellet was resuspended in 0.8 mL cold washing buffer by pipetting up and down 20-30 times. The washed nuclei were centrifuged at 500 X g for 5 min. The supernatant was removed and discarded. The nuclei were ready to proceed into the fixation step.

#### Formaldehyde and DSP fixation

After cell nuclei were isolated and washed, 3.7% formaldehyde (1X PBS and diluted in Nuclease-Free water) and 500 µM DSP (ThermoFisher #PIA35393, 1 mg stock dissolved in 50 µL DMSO and then diluted in 1X PBS ) were directly added to the pellet to fix nuclei, the incubation time was 10 min. For 5 × 10 ^6^-10 ^7^ cells, 5 mL 3.7% formaldehyde and DSP were added. After fixation, the nuclei were incubated with 1X PBS/0.1 %triton for 5 min twice to improve permeabilization. After this, the nuclei were incubated with clampFISH wash buffer/0.25% triton for 20 min to achieve further permeabilization. The nuclei were ready to do clampFISH treatment.

#### clampFISH

The clampFISH procedure was conducted as in previous work ^14^ with several modifications to increase nuclear RNA FISH signal.

We cultured cells on glass coverslips until they reached approximately 70% confluence. The cells were washed twice with PBS and then fixed in 4% formaldehyde in PBS at room temperature for 10 minutes. After aspirating the formaldehyde, we rinsed the cells twice with PBS and stored them in 70% ethanol at 4 °C. For hybridization, we incubated the cells for at least 4 hours at 37 °C in a buffer containing 10% dextran sulfate, 5× SSC, 0.25% Triton-X-100 and 20% formamide, along with 0.5 μL of the primary ClampFISH probe. Probes were designed according to Rouhanifard *et al*. ^14^.

Next, we performed two 30-minute washes at 37°C in wash buffer (2× SSC, 10% formamide). We then incubated the cells for at least 2 hours at 37 °C in a hybridization buffer with 1 μL of the secondary ClampFISH probe, followed by the same wash procedure. After the second wash, we initiated the click chemistry reaction by adding a solution of 75 μM CuSO4 premixed with 150 μM BTTAA ligand (Jena Biosciences) and 2.5 mM freshly prepared sodium ascorbate (Sigma) in 2× SSC. The samples were incubated for 30 minutes at 37 °C and briefly rinsed with wash buffer.

We continued the amplification cycles by alternating between secondary and tertiary ClampFISH probes, followed by the respective wash steps until the desired signal amplification was achieved. After the final wash, the cells were rinsed once with 2× SSC containing DAPI and once with antifade buffer (10 mM Tris, pH 8.0, 2× SSC, 1% w/v glucose). The samples were then mounted for imaging in an antifade buffer containing catalase (Sigma) and glucose oxidase (Sigma) to prevent photobleaching.

For suspension cells, 0.25% Triton X-100 was added to the wash buffer; the cells were stored in 2X SSC/0.25% triton after clampFISH with desired rounds ready for flow cytometry sorting. Sequences for clampFISH probes can be found in **Supplementary Table 1**.

### Flow cytometry and sorting

For nuclampFISH, we sorted on *the EEF2* intron and *NEAT1* based on the nuclampFISH signal using the BECKMAN Sorter (Cell Sorter), which uses 638 nm excitation and 660 nm emission.

### DSP cross-linking reversal

To reverse the DSP crosslinking for chromatin accessibility assays, the sorted cells were treated with 25 mM DTT at 37°C for 30 minutes, according to the manufacturer’s instructions. The cells were then washed with 1X PBS once and ready for the chromatin accessibility assay.

### RNA extraction, RT-qPCR

RNA was extracted using Trizol (Thermo Fisher #15596026) and resuspended in NF water. DNase (Thermo Fisher #AM1907) was used to remove the contaminant DNA. RNA was reverse transcribed (Fisher Scientific #18-080-044). The Luna qPCR (NEB #M3004) was used to quantify the generated cDNA. The OneTaq RT-PCR mix (NEB #E5315) was used to amplify the generated cDNA. Sequences for primers can be found in **Supplementary Table 1**.

### Chromatin Accessibility Assay

After nuclei were sorted out based on the nuclampFISH signal and DSP crosslinking was cleaved, the cell’s chromatin was extracted, and the chromatin accessibility assay (EpiQuik Chromatin Accessibility Assay Kit) was performed according to the manufacture protocol (Epigentek, #P-1047-48). The changes were made so that the chromatin with/without nuclease treatment was quantified by the previous qPCR protocol ^28^, and the housekeeping gene *HPRT1* was set as a control.

### Image acquisition

Microscopy was performed using a Nikon inverted research microscope eclipse Ti2-E/Ti2-E/B using a Plan Apo λ 20X/0.75 objective or Plan Apo λ 100X/1.45 oil objective. The Epi-fi LED illuminator linked to the microscope assured illumination and controlled the respective brightness of four types of LEDs of different wavelengths. Images were acquired using the Neutral Density (ND16) filter for probes coupled with Alexa 488, Alexa 594, Alexa 647, and cy3. Images were acquired and processed using ImageJ. Images acquired using the Neutral Density (ND16) filter are false-colored gray.

### Image analysis and quantification

We initially segmented and thresholded the images using a custom Matlab software suite (rajlabimage tools: https://github.com/arjunrajlaboratory/rajlabimagetools/wiki). Cell segmentation was performed manually by drawing boundaries around non-overlapping cells. The software then fits each detected spot to a two-dimensional Gaussian profile, specifically on the z plane on which it occurs, to ascertain subpixel-resolution spot locations. Colocalization took place in two stages. In the first stage, the software searched for the nearest neighbor for the smFISH probe with the nuclampFISH probe within a 2.5-pixel (360-nm) window to determine colocalization.

## Author Contributions

Y.L. and S.H.R.: conceived and designed the experiments. Y.L. performed the experiments. Y.L., Y.Q. and K.N.: analyzed the data. Y.L. and S.H.R.: wrote the paper with contributions from all authors.

## Acknowledgments

The authors thank the Institute for Chemical Imaging of Living Systems (CILS)-Northeastern University (Core facilities Director: Guoxin Rong) for using and helping with flow cytometry and flow sorting.

## Notes

The authors declare no competing financial interest.

## REFERENCES

1. Raj, A., Peskin, C. S., Tranchina, D., Vargas, D. Y. & Tyagi, S. Stochastic mRNA synthesis in mammalian cells. PLoS Biol. 4, e309 (2006).

2. Zenklusen, D., Larson, D. R. & Singer, R. H. Single-RNA counting reveals alternative modes of gene expression in yeast. Nat. Struct. Mol. Biol. 15, 1263–1271 (2008).

3. Suter, D. M. et al. Mammalian genes are transcribed with widely different bursting kinetics. Science 332, 472–474 (2011).

4. Coleman, R. A. et al. Imaging Transcription: Past, Present, and Future. Cold Spring Harb. Symp. Quant. Biol. 80, 1–8 (2015).

5. Raj, A., Rifkin, S. A., Andersen, E. & van Oudenaarden, A. Variability in gene expression underlies incomplete penetrance. Nature 463, 913–918 (2010).

6. Park, P. J. ChIP-seq: advantages and challenges of a maturing technology. Nat. Rev. Genet. 10, 669–680 (2009).

7. Johnson, D. S., Mortazavi, A., Myers, R. M. & Wold, B. Genome-wide mapping of in vivo protein-DNA interactions. Science 316, 1497–1502 (2007).

8. Wissink, E. M., Vihervaara, A., Tippens, N. D. & Lis, J. T. Nascent RNA analyses: tracking transcription and its regulation. Nat. Rev. Genet. 20, 705–723 (2019).

9. Wheat, J. C. et al. Single-molecule imaging of transcription dynamics in somatic stem cells. Nature 583, 431–436 (2020).

10. Senecal, A. et al. Transcription factors modulate c-Fos transcriptional bursts. Cell Rep. 8, 75–83 (2014).

11. Choi, H. M. T. et al. Third-generation in situ hybridization chain reaction: multiplexed, quantitative, sensitive, versatile, robust. Development 145, (2018).

12. Garcia-Perez, L. et al. A Novel Branched DNA-Based Flowcytometric Method for Single-Cell Characterization of Gene Therapy Products and Expression of Therapeutic Genes. Front. Immunol. 11, 607991 (2020).

13. Kishi, J. Y. et al. SABER amplifies FISH: enhanced multiplexed imaging of RNA and DNA in cells and tissues. Nat. Methods 16, 533–544 (2019).

14. Rouhanifard, S. H. et al. ClampFISH detects individual nucleic acid molecules using click chemistry– based amplification. Nat. Biotechnol. 37, 84–89 (2018).

15. Dardani, I. et al. ClampFISH 2.0 enables rapid, scalable amplified RNA detection in situ. Nat. Methods 19, 1403–1410 (2022).

16. Post-transcriptional splicing can occur in a slow-moving zone around the gene. https://elifesciences.org/reviewed-preprints/91357.

17. Kotake, Y. et al. Splicing factor SF3b as a target of the antitumor natural product pladienolide. Nat. Chem. Biol. 3, 570–575 (2007).

18. Femino, A. M., Fay, F. S., Fogarty, K. & Singer, R. H. Visualization of single RNA transcripts in situ. Science 280, 585–590 (1998).

19. Raj, A., van den Bogaard, P., Rifkin, S. A., van Oudenaarden, A. & Tyagi, S. Imaging individual mRNA molecules using multiple singly labeled probes. Nat. Methods 5, 877–879 (2008).

20. Liu, C.-W. et al. Accurate Measurement of Formaldehyde-Induced DNA–Protein Cross-Links by High-Resolution Orbitrap Mass Spectrometry. Chem. Res. Toxicol. 31, 350–357 (2018).

21. Tayri-Wilk, T. et al. Mass spectrometry reveals the chemistry of formaldehyde cross-linking in structured proteins. Nat. Commun. 11, 3128 (2020).

22. Ongay, S. et al. Cleavable Crosslinkers as Tissue Fixation Reagents for Proteomic Analysis. Chembiochem 19, 736–743 (2018).

23. Tsompana, M. & Buck, M. J. Chromatin accessibility: a window into the genome. Epigenetics Chromatin 7, 33 (2014).

24. Mansisidor, A. R. & Risca, V. I. Chromatin accessibility: methods, mechanisms, and biological insights. Nucleus 13, 236–276 (2022).

25. Berrocal, A., Lammers, N. C., Garcia, H. G. & Eisen, M. B. Kinetic sculpting of the seven stripes of the Drosophila even-skipped gene. Elife 9, (2020).

26. Pandya-Jones, A. & Black, D. L. Co-transcriptional splicing of constitutive and alternative exons. RNA 15, 1896–1908 (2009).

27. Schwarzkopf, M. et al. Hybridization chain reaction enables a unified approach to multiplexed, quantitative, high-resolution immunohistochemistry and in situ hybridization. Development 148, (2021).

28. Liu, Y. et al. Paired Capture and FISH Detection of Individual Virions Enable Cell-Free Determination of Infectious Titers. ACS Sens 8, 2563–2571 (2023).

