## Supporting Information for "NuclampFISH enables cell sorting based on nuclear RNA expression for chromatin analysis"

*DSP reversal crosslinking is compatible with chromatin accessibility assay.*

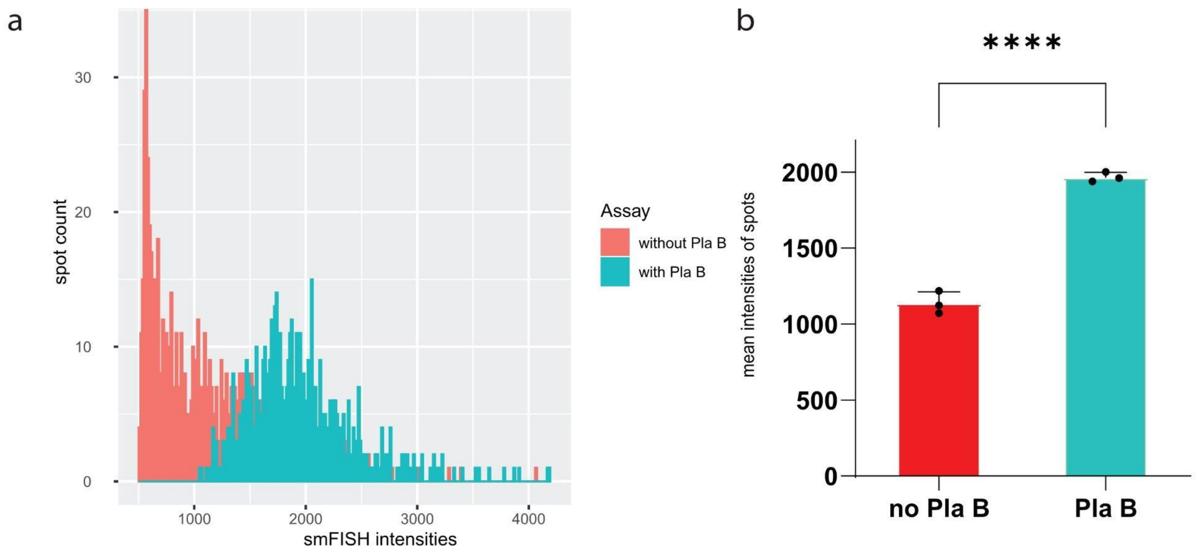

**Supplementary Figure 1:** smFISH signal of *EEF2* intron increases after Pla B treatment. a. Histogram of single spots intensities of smFISH with Pla B and without Pla B treatment in Figure 2a. b. Bar graph of mean intensities of smFISH spots with Pla B and without Pla B treatment. n = 3 biological replicates (bars represent mean ± SD).*p < 0.05, **p < 0.01, ***p < 0.001, ****p < 0.0001.

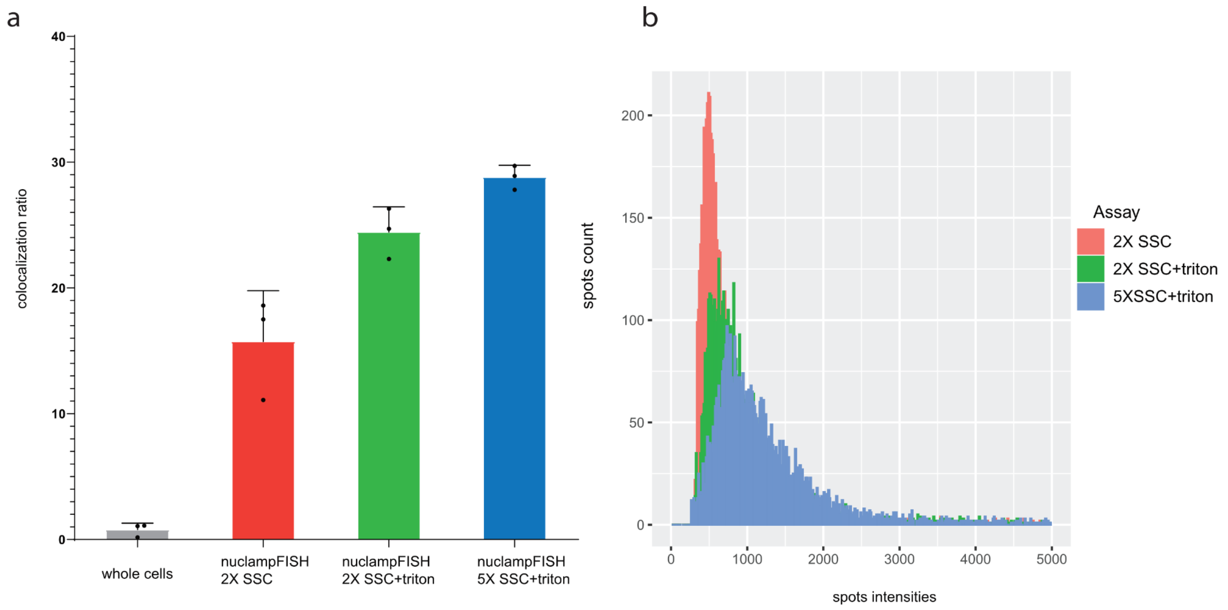

**Supplementary Figure 2:** Optimized conditions of nuclampFISH with triton and increased SSC concentration. a. Improvement of colocalization ratio of nuclampFISH after nucleus isolation, adding triton and increasing SSC concentration. b. Histogram of single spots intensities after treatment of triton and increased SSC concentration. n = 3 biological replicates (bars represent mean ± SD).

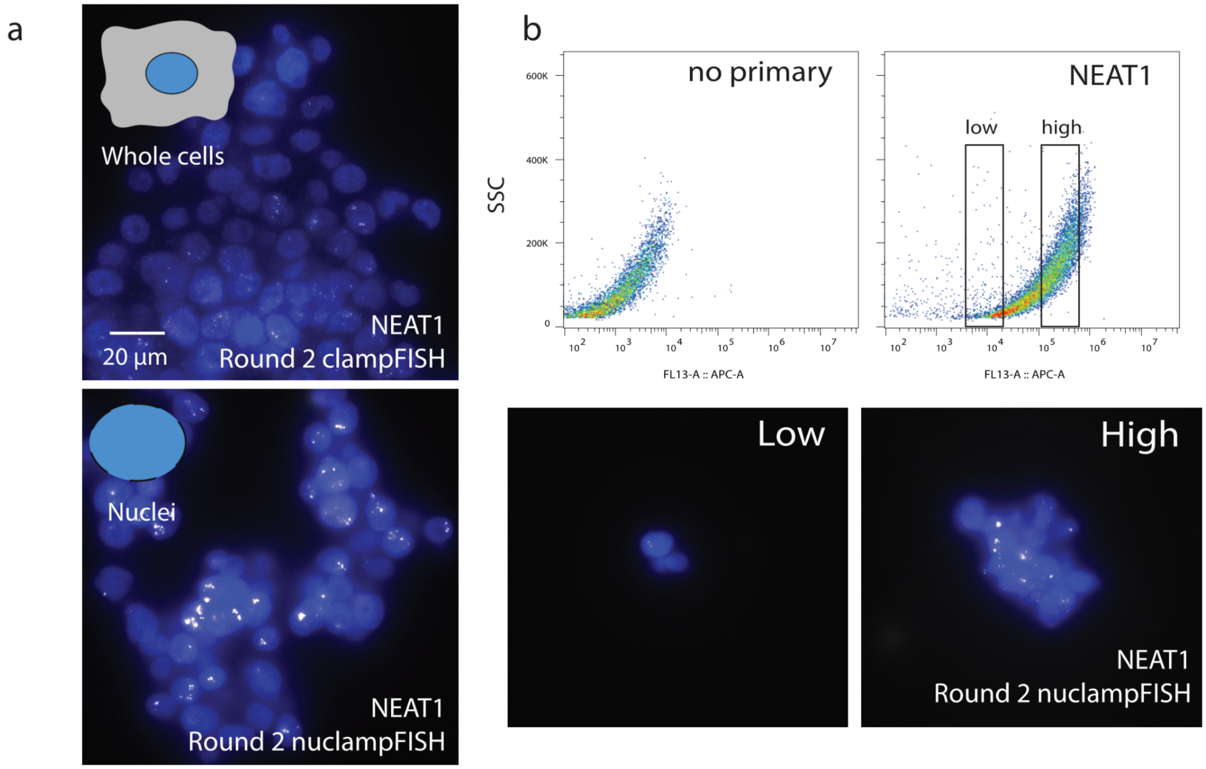

**Supplementary Figure 3:** NuclampFISH for long-non coding RNA *NEAT1*. a. ClampFISH images and nuclampFISH images of *NEAT1*. b. Flow cytometry data of nuclampFISH, dot plot of nuclampFISH signal compared to no primary negative control group, based on the low and high signal in the flow cytometry, sorted cells were imaged.

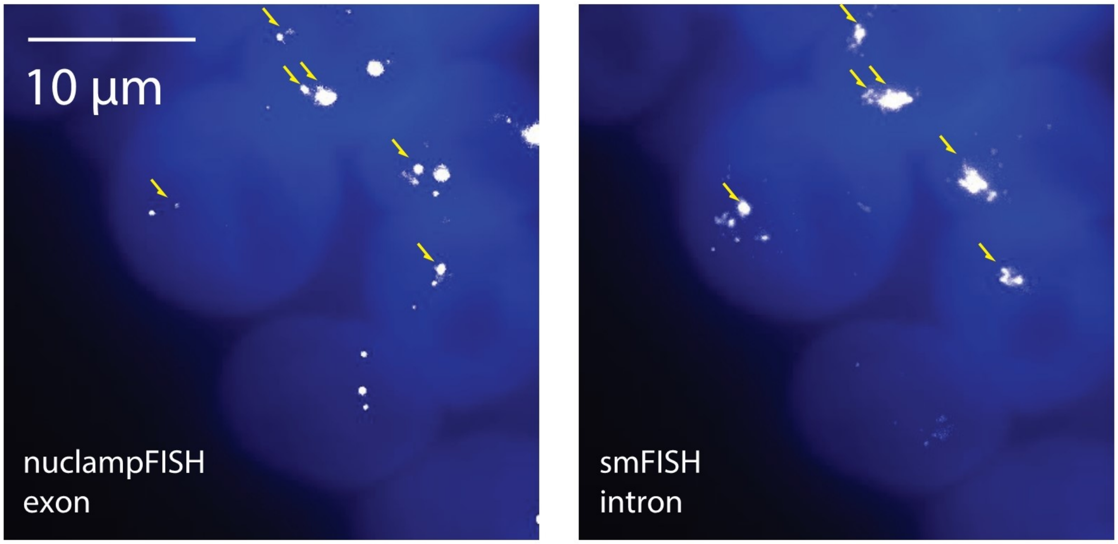

**Supplementary Figure 4:** NuclampFISH for *TMSF1* genes. NuclampFISH images of *TMSF1* exon were overlapped with smFISH images of *TMSF1* intron. Arrow indicates the transcription sites overlaid in both images.

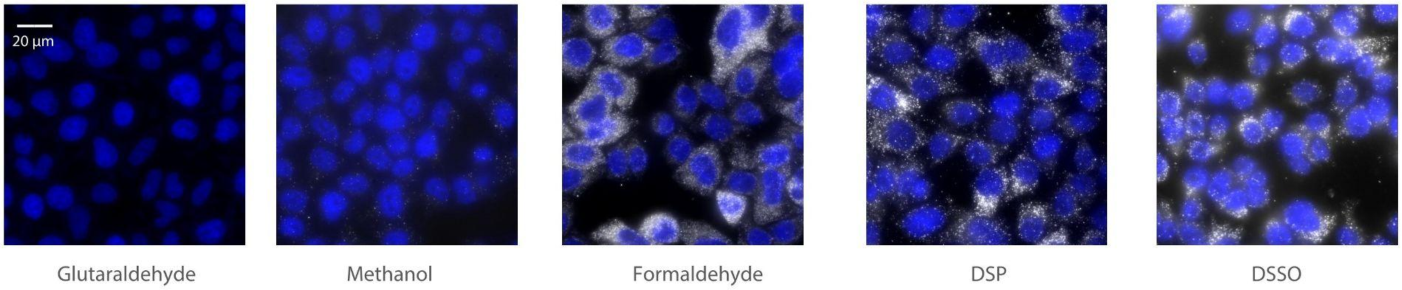

**Supplementary Figure 5:** Compatibility of different crosslinkers for clampFISH. Images of different fixation reagents of clampFISH for *HIST1H4E* gene.

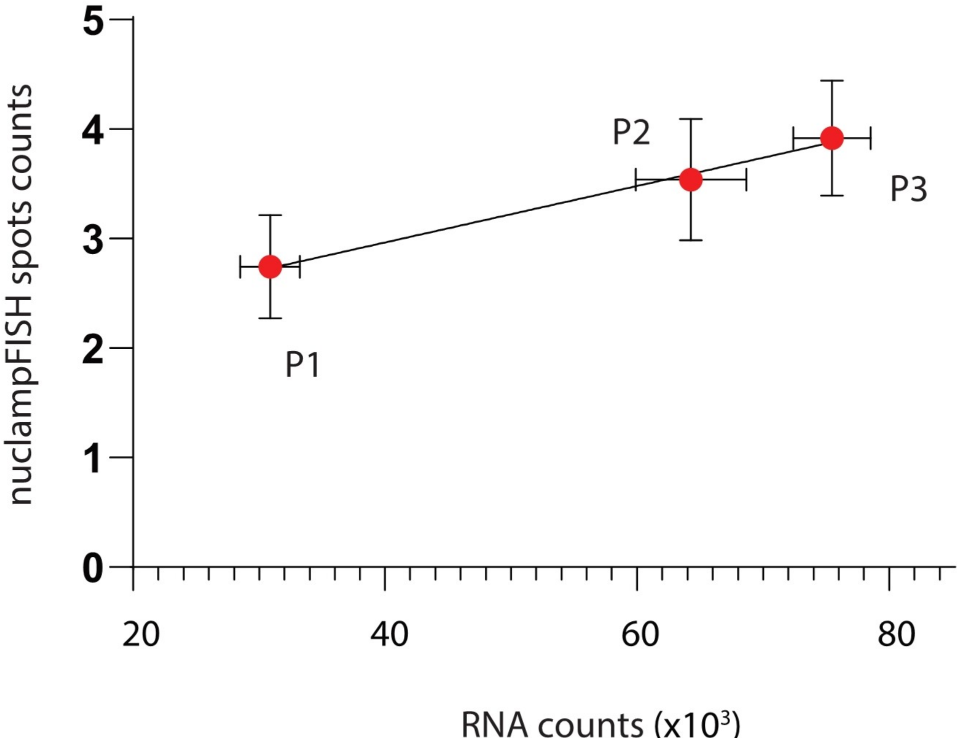

**Supplementary Figure 6:** Correlation of RT-pPCR counts with nuclampFISH spots counts after flow sorting. n=3 biological replicates (mean ± SD).

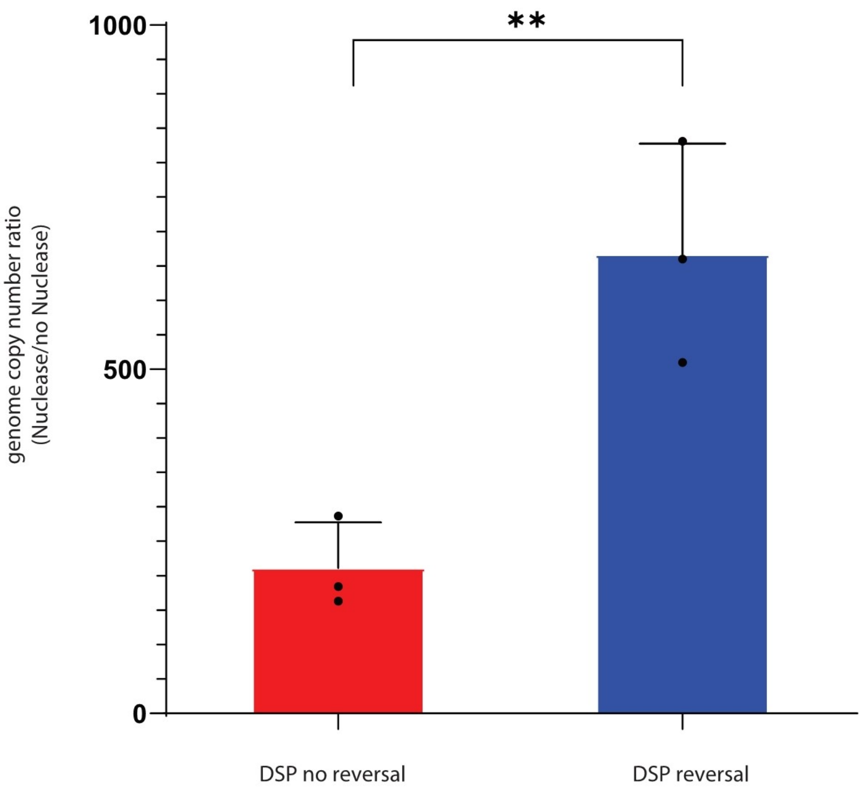

**Supplementary Figure 7:** DSP reversal crosslinking is compatible with chromatin accessibility assay. Quantitative comparison of chromatin accessibility between DSP without reversal crosslinking and DSP with reversal crosslinking. **p < 0.01
